# Some generic measures of the extent of chemical disequilibrium applied to living and abiotic systems

**DOI:** 10.1101/327783

**Authors:** B. F. Intoy, J. W. Halley

## Abstract

We report results of evaluation of several measures of chemical disequilibrium in living and abiotic systems. The previously defined measures include *R*_*T*_ and *R*_*L*_ which are Euclidean distances of a coarse grained polymer length distribution from two different chemical equilibrium states associated with equilibration to an external temperature bath and with isolated equilibration to a distribution determined by the bond energy of the system, respectively. The determination uses a simplified model of the energetics of the constituent molecules introduced earlier. We evaluated the measures for data from the ribosome of E. Coli, a variety of yeast, the proteomes (with certain assumptions) of a large family of prokaryotes, for mass spectrometric data from the atmosphere the Saturn satellite Titan and for commercial copolymers. We find with surprising consistency that *R*_*L*_ is much smaller than *R*_*T*_ for all these systems. Small *R*_*L*_ may be characteristic of systems in the biosphere.

## I. INTRODUCTION

To define the mission of the search for extraterrestrial biology with quantitative precision, it would be desirable to specify measures of molecular systems which are more generic than searches for particular molecules or combinations of molecules, such as those in the earth’s biosphere. That is because it is not known that all nonequilibrium systems having lifelike characteristics will turn out to be at all chemically similar to the terrestrial biosphere. Designing such measures is a goal of our program of study of Kauffman-like models of prebiotic evolution. The choice of such measures constitutes an implicit definition of the meaning of the term ‘lifelike’. However if the measures are quantitatively well defined, then one can use them without engaging in a debate about whether certain ranges of their values imply, necessarily or sufficiently, a ‘lifelike’ system.

Here we evaluate several such generic measures for a set of experimentally known systems, including some which are in the biosphere and others which are not. A goal is to determine how well the measures distinguish living from nonliving systems and what they reveal about the differences. The measures used here were first applied by us to the results of simulations of a Kauffman-like model as described in [1]. However we did not need to assume a detailed correspondence between that model and the experimentally observed systems described here in order to carry out the program reported in this paper. We do need to assume that the systems contain only linear polymers and we make some simplifying assumptions about the bonding energies as described in more detail below. In particular, the assumptions about the energy allot an upper bound to the energy, and thus allow equilibrium with negative temperatures as discussed below. These assumptions are well satisfied for some of the systems under study and for others they are more problematical as we will discuss.

The main experimental inputs to the analysis presented here are the molecular weight density distributions, the numbers of monomers available for forming polymers (parameter *b* defined below) and two parameters characterizing the size of a polymer coil of L monomers. The main outputs include values of two dimensionless parameters termed *R*_*T*_ and *R*_*L*_ which describe how far the observed system is from chemical equilibrium with an external thermal bath and from chemical equilibrium if isolated, respectively. Detailed definitions are provided in section II.

The systems we study are the proteomes of 4,555 prokaryotes, the ribosomes in E. coli, a budding yeast, five artificial copolymers, and the atmosphere of Titan. The choice of systems was made in order to provide a sampling of living systems and some engineered system (the copolymers). Titan’s atmosphere was selected because Titan’s atmosphere is reported [2] to contain an abundance of organic molecules and may be characteristic of a prebiotic environment [4].

We find some interesting trends: We find that all the systems are closer to isolated equilibrium with a high temperature which is negative for the copolymer and Titan systems than they are to chemical equilibrium with the (positive) ambient temperature of their surroundings. The living systems have a sharply defined *R*_*T*_ value different from the values found for the other systems. It appears that a small *R*_*L*_ and a large well defined *R*_*T*_ distinguish the living from the nonliving systems. This feature is closely related to the fact that in peptide systems under biological conditions the bonding energy as defined in our model is negative whereas it is positive for the other systems. We present detailed molecular weight distributions to contrast what is observed experimentally with the corresponding equilibria.

In the next section we define the essential features and assumptions which allow definition and evaluation of the measures *R*_*L*_, *R*_*T*_ and other generic characteristics. We focus first on the case in which all the bond energies are the same followed by a brief discussion of the case in which there is more than one bond energy. Section III reports the processes by which we extracted the needed information from the raw experimental data in each of the cases considered. In section IV we report the results and section V contains a discussion and conclusions.

## II. ANALYSIS

We first describe the case in which all the polymeric bond energies are the same, as we have assumed for all systems except the Titan atmosphere. Polymers are assumed to consist of strings of monomers with *b* types possible for each monomer. We took *b* = 20 for the ribosomes in E. coli, the prokaryotes and for yeast and *b* = 2 for the Titan data and the copolymer data. To any polymer of length *L* we attribute an energy −(*L* − 1)Δ where Δ is a real number which is the bonding energy between two monomers. It is taken to be positive for the copolymers and the Titan atmosphere but negative for the peptide bonds in the biological systems as discussed in the next section. The total energy *E* of any population {*n*_*m*_} of polymers in which *n*_*m*_ is the number of polymers of type *m* is 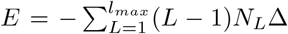 Here the 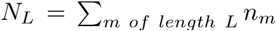 is the same set of macrovariables used in [5] and [6]. We denote the total number of polymers *N* in a sample by is 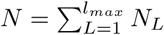 However, in contrast to the situation in the dynamic simulations described in [1], the input data for calculation of equilibrium distributions are not *N* and *E* but the volumetric polymer concentration *ρ* = *N/V* where *V* is the solution volume and the volumetric energy density *e* = *E/V*. To take entropic account of the dilution of the experimental sample we introduce a microscopic length where *R*_0_ *L*^*v*^ is a length related to the polymer persistence length and *ν* is an index which would be 1*/*2 for a random walk. We report the numerical values for *ν* and *R*_0_ used for the various systems considered in the next section. We modify the expression for the entropy used in [1] to take account of the number of ways to distribute *N* polymers in a volume *V* as follows: *S/k*_*B*_ = ln *W* with

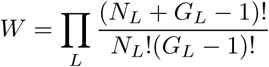

and 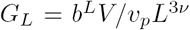 and 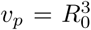. *b* is the number of monomers available for inclusion in the polymers in the system and *ν* is a dimensionless number which is 1/2 for a random walk. The expression is identical to the one used in [1] except for the factor 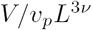 in the degeneracy *G*_*L*_. Note that a similar configurational entropy factor was used by us in an earlier paper [6]. With this modification we apply Stirling’s approximation and maximize the entropy subject to the density and energy constraints associated with the experimental data discussed in the next section. We have

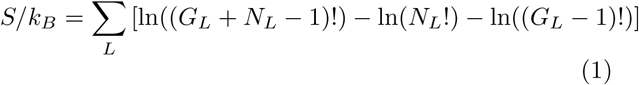

Proceeding in the standard way to maximize the entropy under these constraints we have when both energy density *e* = *E/V* and polymer number density *ρ* = *N/V* are fixed that the values 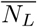 of the populations which maximize this entropy are

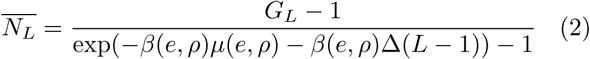

Here the parameters *β*(*e, ρ*) and *μ*(*e, ρ*) are determined from the total energy density *e* and polymer number density *ρ* = *N/V* by the implicit equations (with (2))

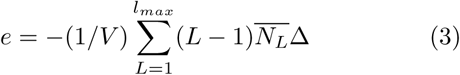

and

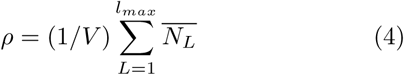

We use the definition of *G*_*L*_ and define 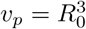 to write these relations as

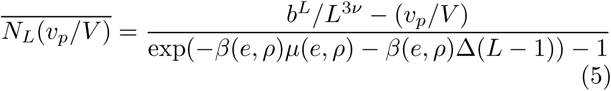

and

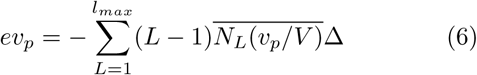

and

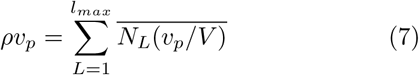

These are in dimensionless form, convenient for solving for *β*(*e, ρ*)*μ*(*e, ρ*) and *β*(*e, ρ*)Δ numerically because they do not involve macroscopically large numbers. The term *v*_*p*_*/V* on the right hand side of (5) is in all cases much less than *b*^*L*^*/L*^3*v*^ and is dropped in the numerical analysis. As before [1] we refer to this equilbrium as ‘isolated’. There is no reference to an external temperature bath. Also, as in [1] we also determine a ‘thermal’ equilibrium distribution by solving (7) with a fixed value of *β* using reported approximate values of the ambient temperature in the experiments considered and making no use of *e*. In each case we can then use (5) to evaluate the polymer length density distributions expected in those two equilibrium states.

Because the systems of interest are not necessarily (and in fact are found not to be) in either kind of chemical equilibrium we used the experimentally observed values of the quantities *N*_*L*_(*v*_*p*_*/V*) to evaluate Euclidean distance in the space of values of sets {*N*_*L*_*v*_*p*_*/V*} between the actual population set {*N*_*L*_*v*_*p*_*/V*} and the ones corresponding to the two kinds of equilibria given by (5) with *β*(*e, ρ*), *μ*(*e, ρ*) in the isolated case and with a fixed ambient *β* and *μ*(*ρ*) in the case in which the system is equilibrated to an external bath. Thus we define two Euclidean distances *R*_*L*_ and *R*_*T*_ in the *l*_*max*_ dimensional space of sets {*N*_*L*_(*v*_*p*_*/V*)} which characterize how far the system of interest is from the two kinds of equilbria:

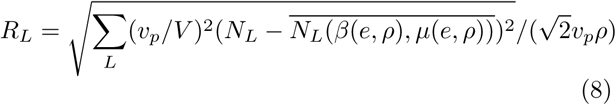

for distance from the locally equilibrated state and

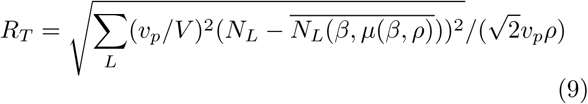

for distance from the thermally equilibrated state.

In the cases of prokaryotes, E. coli ribosomes, yeast and the copolymers which we evaluate, we use the assumptions as just described. However in the analysis of the data from the Titan atmosphere, we take account of the large difference between the energies of CC and CN bonds and NN bonds by reformulating the description with two bond strengths as described in Appendix A and reference [9]. We show there that an accurate treatment of that case gives results very close to those obtained by using the same model as the one described above, but with an average bond strength 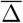 of

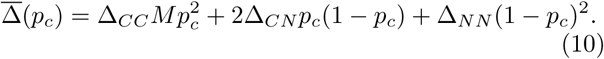

*p*_*c*_ is the atomic fraction of the atmosphere which is carbon. (We assume Δ_*CC*_ = Δ_*CN*_ as discussed further in the next section and Appendix B).

## III. EXTRACTION OF POPULATION DISTRIBUTIONS FROM DATA

Here we describe how data was extracted from data available on a budding yeast, 4,555 prokaryotes, the ribosomes in E. Coli, the commercial copolymers and mass spectrographic data on the atmosphere of Titan in order to compute the characteristic parameters *R*_*L*_, *R*_*T*_, *βμ* and *β*Δ defined in the preceding section. We report a comparison of the results in the next section in order to determine the extent to which these parameters may distinguish living from lifelike or nonliving systems.

### A.Yeast

We used data on protein population distributions in the budding yeast *Saccharomyces cerevisiae* from reference [16] with[7] *R*_0_ = 1.927Å and *v* = 0.588 in solving equations (6) and (7). For determining the chemical potential in the case of equilibrium with an external temperature bath (equation (7)) we used a temperature of 293 K and a bond energy [8] of −2.2 kcal/mol. It is important to note that for this and the other protein systems Δ is taken to be negative, meaning that it costs energy to make a peptide bond. The value of −2.2 kcal/mol taken from reference [8] is the Gibbs free energy of reaction which can be seen from the description of the model in [1] to be the appropriate energy to identify with Δ within our model.

### B. Ribosomes in E. coli

We used protein distribution data from [20]. There are approximately 50000 ribosomes per cubic micron in an E. coli cell [21] the ribosome giving a value for *Nv*_*p*_*/V* of about 1.86 ×10^−5^ assuming that, in equilibrium, the proteins would be denatured and evenly distributed throughout the cell. We took *R*_0_ = 1.927Å and *v* = 0.588 as for the yeast and prokaryotes.

### C. Prokaryote Proteins

We used data from the Kyoto Encyclopedia of Genes and Genomes[14] (KEGG) (the faa, fasta amino acid file) to extract approximate length distributions for 4,555 prokaryotes assuming that there is only one of each protein in each prokaryotic cell. The KEGG data gave amino acid sequences for all the proteins, from which the coarse grained number *N*_*L*_ of all proteins of each length *L* was extracted. To get a volumetric distribution *N*_*L*_*/V* by averaging data in [15] we obtained an approximate average volumetric protein density of 2.5 million proteins per cubic micron which was used for all the prokaryotes when solving equations 7 and 6. As for yeast, the form[7] 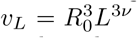 with *R*_0_ = 1.927Å and *v* = 0.588 was used in the solutions to equations 7 and 6. For thermal equilibrium calculations a temperature of 20 degrees Celsius and a bond energy of −2.2 kcal/mol was used.

### D. Copolymers

We used data from references [17, 18]. The experimental details are summarized in Table I. The data references given in Table I give the relative abundances of polymers as a function of their composition. The table also mentions the monomers used in the systems which were either polystyrene-polyisoprene or polystyrene polybutadiene. For example the data from [17] gives the relative abundances *f* (*l*_*s*_, *l*_*i*_) of polymers in a solution of a polystyrene-polyisoprene system as a function of their styrene *l*_*s*_ and isoprene *l*_*i*_ compositions, as extracted from mass spectrometry data. To get values of *f*_*L*_ we sum all the values of *f* (*l*_*s*_, *l*_*i*_) for which *l*_*s*_ +*l*_*i*_ = *L*. From [17], the total number of monomer blocks in the copolymer system was 8.6 mol, consisting of 0.7 mol of styrene and 7.9 mol of isoprene. From the length distribution {*f*_*L*_}the average polymer length is calculated to be approximately 28.9. Dividing the total number of monomer blocks by the average polymer length gives the total number of polymers to be approximately *N* = 1.79 ×19^23^. The experiment took place in a *V* = 6 Liter reactor vessel giving *N/V*. For solving equations (6) and (7) we obtained *v*_*p*_ for this system using listed values for the persistence lengths *l*_*p*_, statistical segment lengths 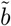 and characteristic ratios *C*_∞_ cited in [25] to obtain the value 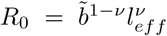 with 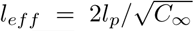 the effective bond length as defined in [25]. For the copolymers we assumed that *ν* = 0.5 and used the average of the *R*_0_ values calculated from the parameters *b, l*_*p*_ and *C*_∞_ for Isoprene and Styrene in [25]. For calculations of the chemical potential *μ* from (7) when the system is in thermal equilibrium with an external thermal bath, a bath temperature of 293 K and the average Carbon-Carbon bond energy of 347 kJ/mol was used [19].

**TABLE I:** Parameters used for the five copolymer systems considered here. *N*_mon_ is the total number of monomer blocks in units of mols, ⟨*L*⟩ is the average polymer length, and *V* is the volume in liters. *R*_0_ values are calculated by taking the average of the two monomer types from [25].

| Data Reference | Monomer types | Figure in this paper | $N_{\text{mon}}$ (mols) | $\langle L \rangle$ | $V$ (Liters) | $R_0$ (Å) | Density ($Nv_p/V$) |
| --- | --- | --- | --- | --- | --- | --- | --- |
| Figure 7 of [17] | Isoprene and Styrene | 8 | 8.6 | 28.9 | 6 | 5.10 | 0.00395 |
| Figure 6-13b of [18] | Isoprene and Styrene | 9 | 1.21 | 32.9 | 2 | 5.10 | 0.00147 |
| Figure 6-13c of [18] | Isoprene and Styrene | 10 | 1.21 | 41.3 | 2 | 5.10 | 0.00117 |
| Figure 6-13d of [18] | Isoprene and Styrene | 11 | 1.21 | 49.9 | 2 | 5.10 | 0.000968 |
| Figure 6-15 Left of [18] | Styrene and Butadiene | 12 | 0.966 | 36.6 | 2 | 5.27 | 0.00116 |

### E. Titan Data Extraction and Analysis

We used mass spectrographic data on the atmosphere of Titan taken by the *Cassini* spacecraft as reported in [2]. The data on mass distributions is reported in [2] in units of charge detected per second. In this paper we only report analysis of the Cassini mass spectroscopy data on detected negative ions. The mass distribution of negative ions contains the widest mass distribution reported, including molecules as massive a 10^4^ daltons. Data are available for detected neutral and positively charged molecules [3] and may be analysed later. We assume in our analysis that all the molecules detected had unit charge (in units of the magnitude of the electron charge). Masses were reported in daltons and converted approximately to monomer units by dividing by an assumed average monomer mass of 13 daltons because the monomers are believed to be predominantly single nitrogen or carbon entities. (We are neglecting the contribution of the hydrogen masses). We converted the data reported in [2] in log scale bins to linear scale bins as described in Appendix A. To extract volumetric densities *N/V* we used the kinetic relation 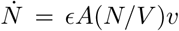 where 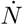is the rate at which particles are detected per per second, *ε* = 0.05 is the detector efficiency [2], *A* is the detector area, reported to be 0.33 *cm*^2^ [2] and *v* is the velocity of the spacecraft relative to the Titan atmosphere for which we used *v* = 6.3*km/sec* from reference [10]. This procedure gave volume densities for data taken at different altitudes above the Titan surface as summarized in Table III. We assumed that the species detected in the mass spectrometer were singly charged. Uncertainties resulting from this procedure are discussed in [2]. For the value of 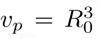 in equations (6) and (7) we used the *R*_0_ = 4.97 Å extracted from data for polyethylene in [25] in the same way that it was done for the copolymers as explained above (with *ν* = 0.5).

**TABLE II:** The average bond energies for carbon and nitrogen [19].

| Bond | Average Bond Energy (kJ/mol) |
| --- | --- |
| C-C | 347 |
| C-N | 305 |
| N-N | 160 |

**TABLE III:** Titan atmospheric densities for various altitudes as detected during the 40th Titan encounter of Cassini which occurred on 05 January 2008.

| Altitude (km) | Figure in this paper | Density ($Nv_p/V$) |
| --- | --- | --- |
| 1013 | 13 | $4.56 \times 10^{-20}$ |
| 1032 | 14 | $3.79 \times 10^{-20}$ |
| 1078 | 15 | $1.72 \times 10^{-20}$ |
| 1148 | 16 | $6.54 \times 10^{-21}$ |
| 1244 | 17 | $7.89 \times 10^{-22}$ |

For determining the equilibrium distribution in the presence of an ambient thermal bath using equation (7) we used an ambient temperature of 120 degrees Kelvin [12].

In applying equations (7) and (6) to the Titan data, we are assuming that all the molecules detected are linear chains, which is certainly not expected to be true [4]. Also, CC and CN bond energies are quite closely similar as described in Table II, but the N-N bond is much weaker. In principle, one should therefore take account of the different bond energies in the model used to obtain equations (6) and (7) which we solve to determine the equilibrium distributions. We have done that, with results described elsewhere [9]. However it turned out that a simplifying ‘mean-field’ approximation works quite well to describe the results and we use that here: We used an ‘average’ bond energy 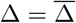 where

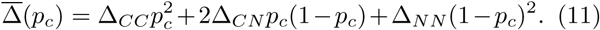

with Δ_*CC*_ = Δ_*CN*_ = 325*kJ/mol* and Δ_*NN*_ = 160*kJ/mol*. *p*_*c*_ is the atomic fraction of the atmosphere which is carbon. We use the value *p*_*c*_ = 0.02 [13]. As explained in Appendix B, we also obtained results using the maximum and minimum values which the bond strength could have at the observed *p*_*c*_ and found that the results were quite insensitive to the change and consistent with the results of [9].

## IV.RESULTS

We summarize the data found for *R*_*T*_ and *R*_*L*_ in Figure 1 with a second view focussing on the region where the data cluster in Figure 2.

**FIG. 1:**
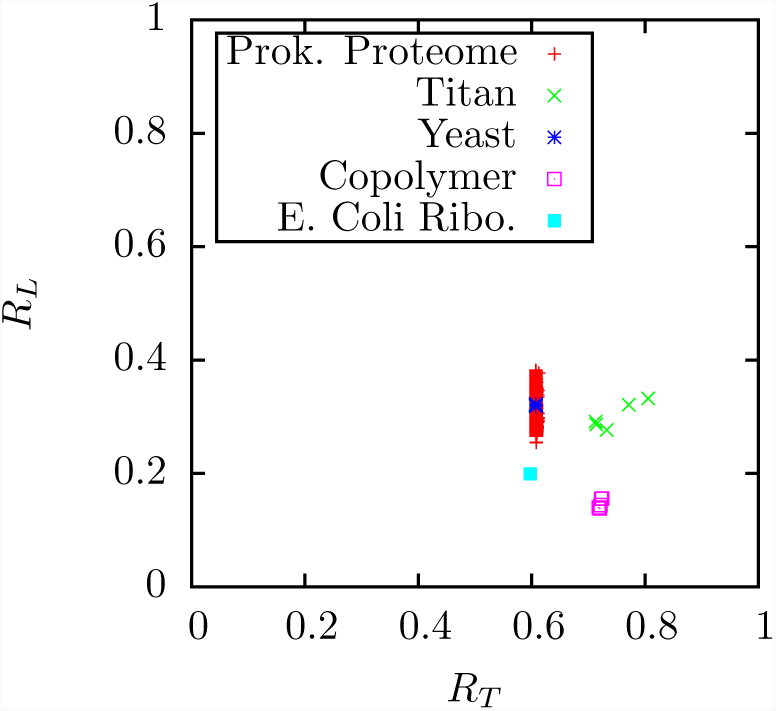
*R*_*T*_ and *R*_*L*_summary scatter plot for all the systems considered.

**FIG. 2:**
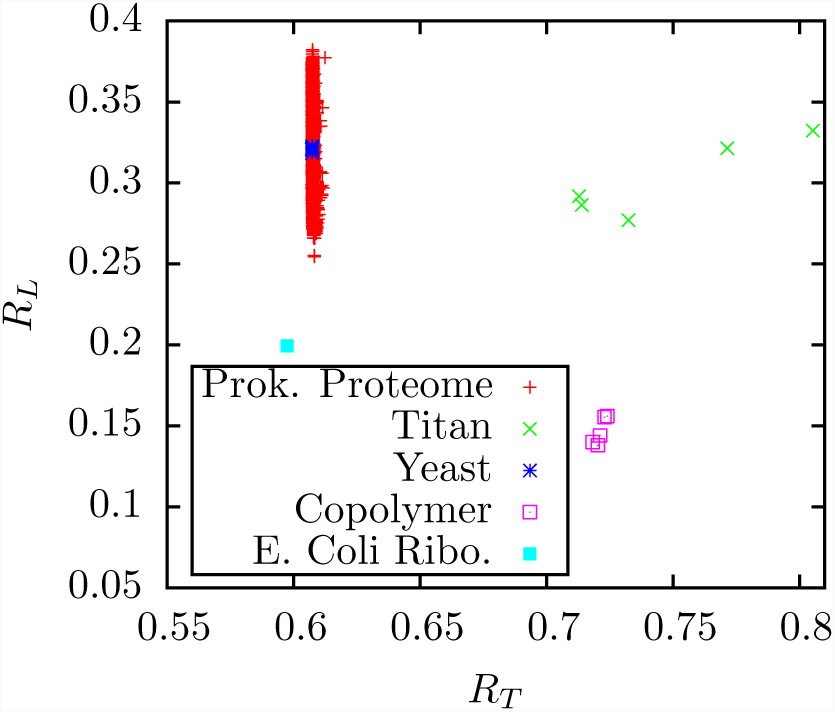
Data from the preceding figure (Fig. 1) showing only the region where system points appear.

The yeast distribution is shown in Figure 3. The distributions for three of the 4,555 prokaryotes analysed are shown in Figures 4-6 and the E. coli ribosomes length distribution appears in Figure 7. Length distributions for five copolymer systems are shown in Figures 8-12. Figures 13-17 compare the observed length distributions with the calculated equilibrium for the Titan data at five elevations. Values of *R*_*L*_ and *R*_*T*_ for the data exhibited in the figures appears in Table IV.

**FIG. 3:**
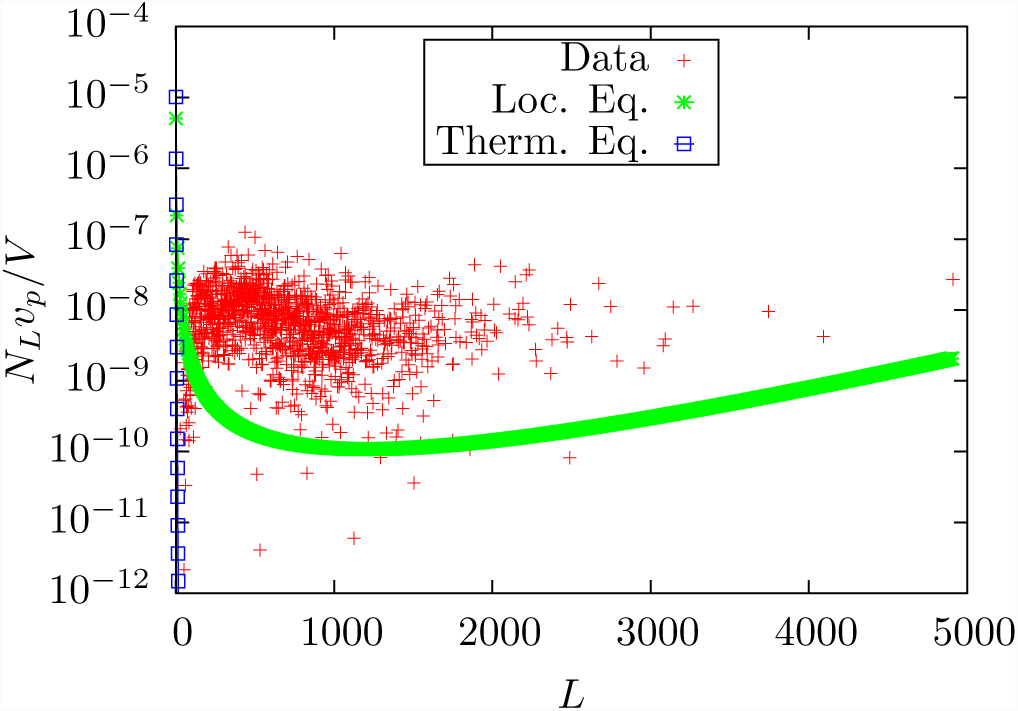
*N*_*L*_*v*_*p*_*/V* values for yeast with glucose as its carbon source together with the corresponding local and thermal equilibrium distributions which are respectively euclidean distances (see equations (8) and (9)) *R*_*L*_ = 0.32 and *R*_*T*_ = 0.61 away from the observed distribution.

**FIG. 4:**
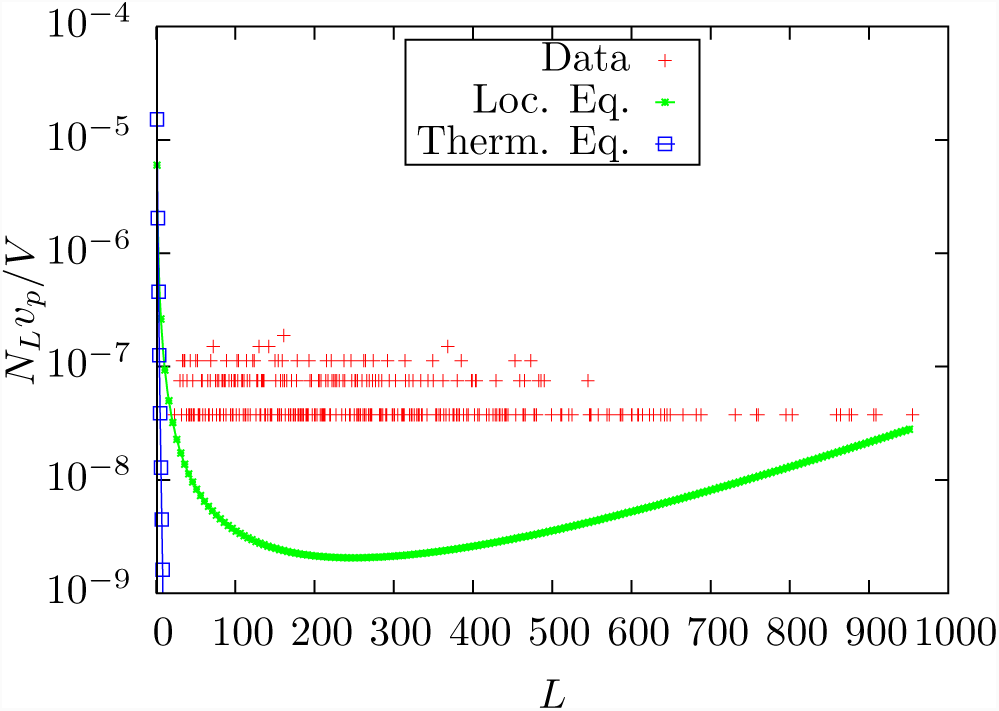
*N*_*L*_*v*_*P*_*/V* values for the proteins found in the prokaryote Buchnera aphidicola JF98 an endosymbiont of Acyrthosiphon pisum which has organism code baw on KEGG and corresponding equilibrium distributions. *R*_*L*_ = 0.25 and *R*_*T*_ = 0.61.

**FIG. 5:**
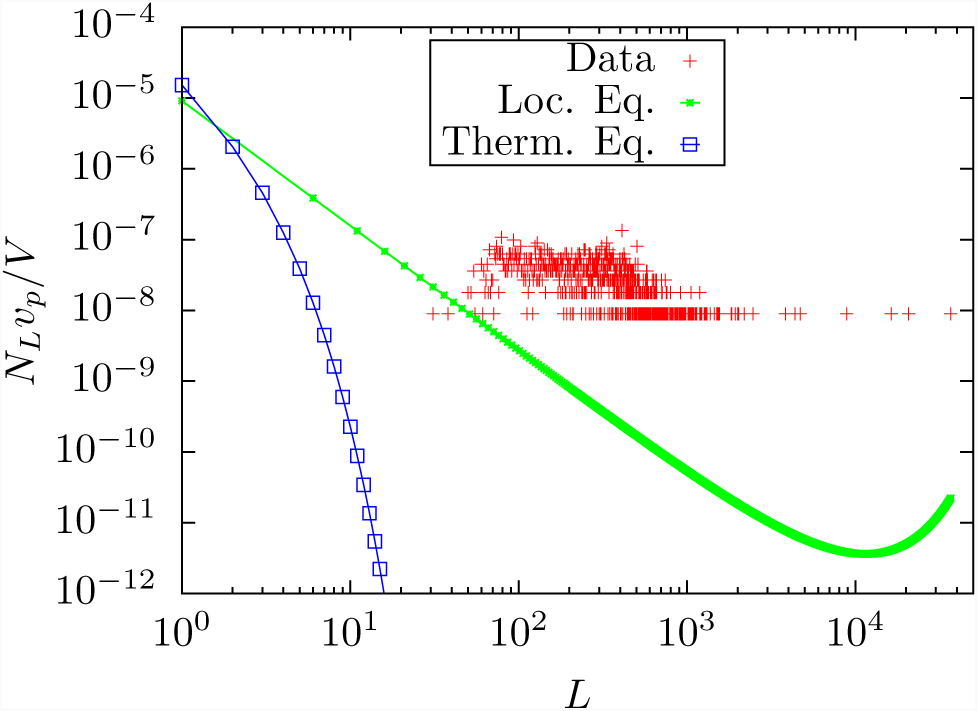
*N*_*L*_*v*_*P*_*/V* values for the proteins found in the prokaryote Chlorobium chlorochromatii CaD3 which has organism code cch on KEGG and and corresponding equilibrium distributions.. *R*_*L*_ = 0.38 and *R*_*T*_ = 0.61.

**FIG. 6:**
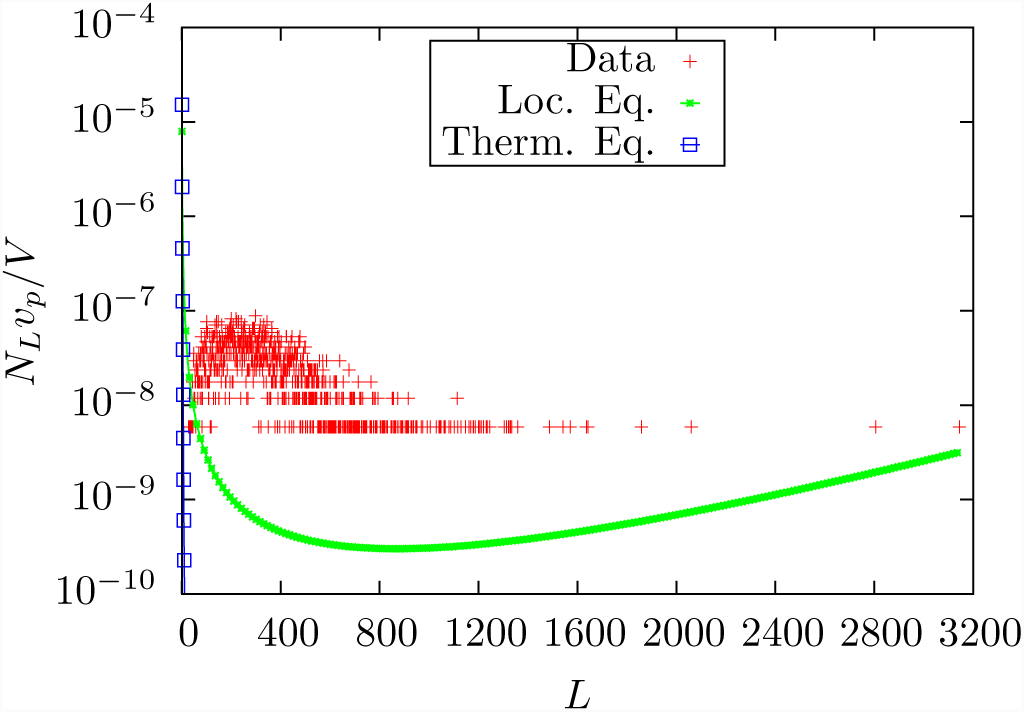
*N*_*L*_*v*_*P*_*/V* values for the proteins found in the prokaryote Corynebacterium variable DSM 44702 which has organism code cva on KEGG and corresponding equilibrium distributions. *R*_*L*_ = 0.33 and *R*_*T*_ = 0.61.

**FIG. 7:**
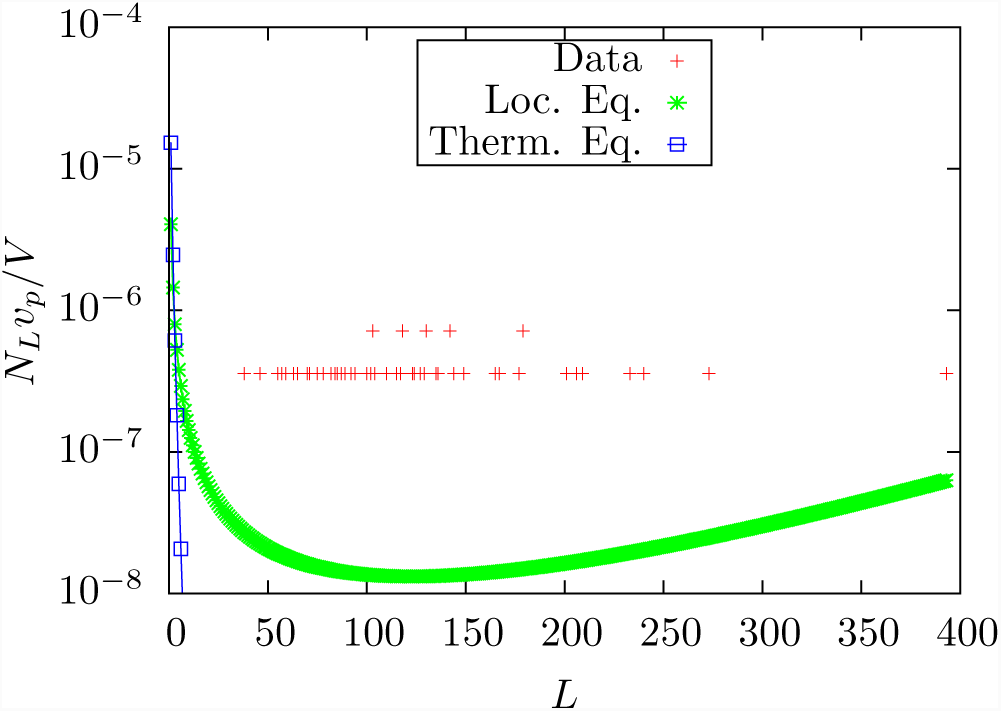
*N*_*L*_*v*_*P*_*/V* values for the proteins in the E Coli ribosome and corresponding equilibrium distributions. *R_L_* = 0.20 and *R_T_* = 0.60.

**FIG. 8:**
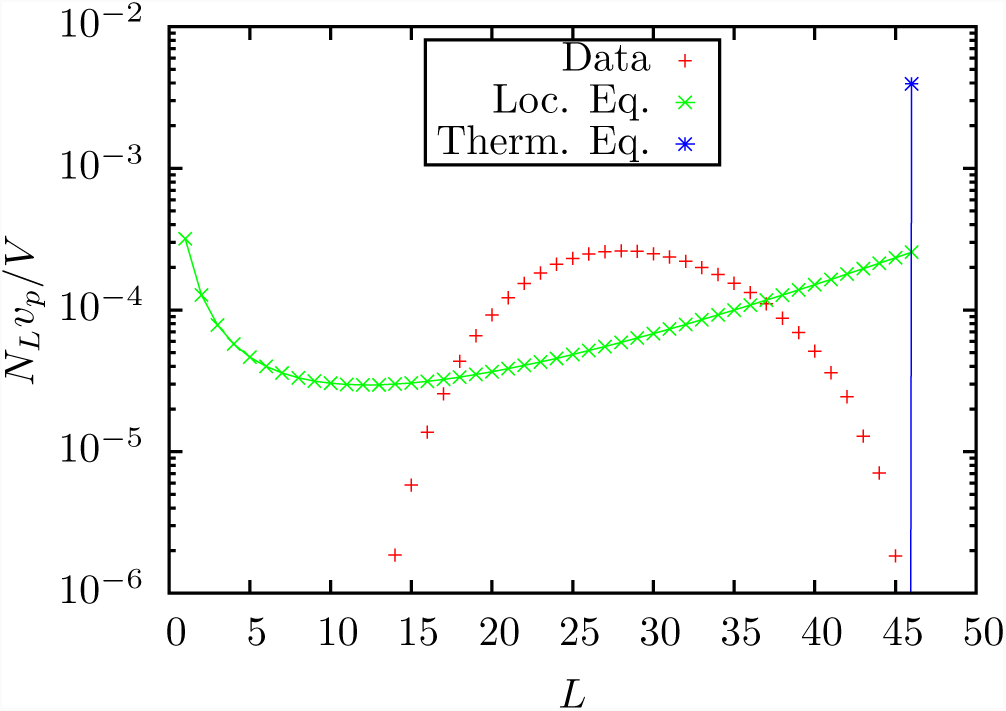
*N*_*L*_*v*_*P*_ */V* values for the data from Figure 7 of [17] and corresponding equilibrium distributions‘. *R*_*L*_ = 0.16 and *R*_*T*_ = 0.72.

**FIG. 9:**
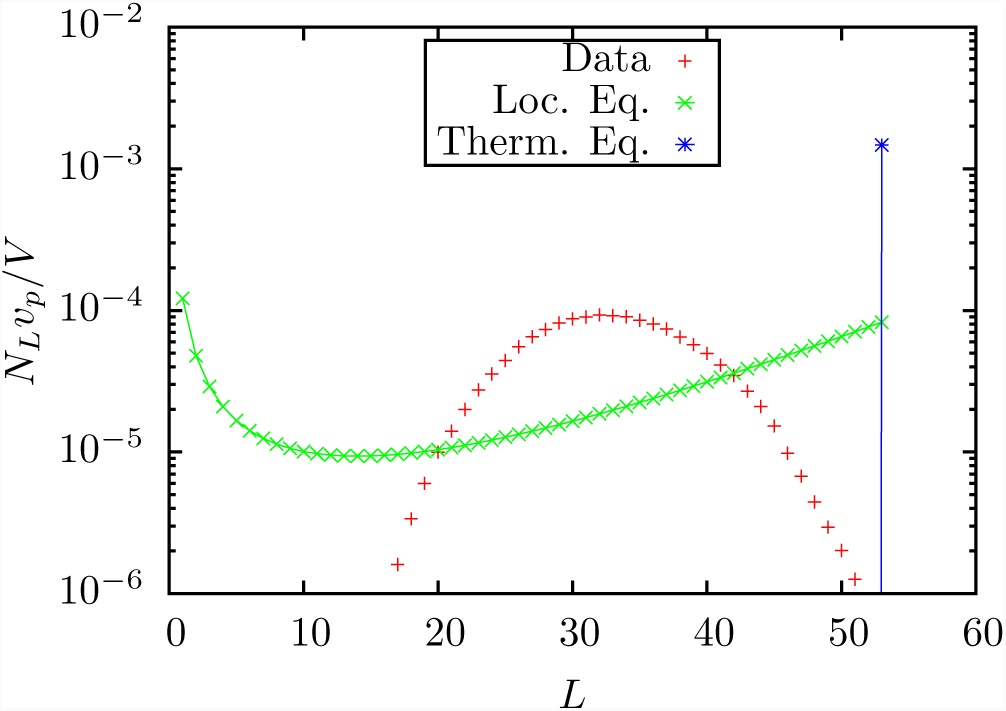
*N*_*L*_*v*_*p*_*/V* values for the copolymer system for which taken data are given in Figure 6-13b of [18] and corresponding equilibrium distributions.. *R*_*L*_ = 0.16 and *R*_*T*_ = 0.72.

**FIG. 10:**
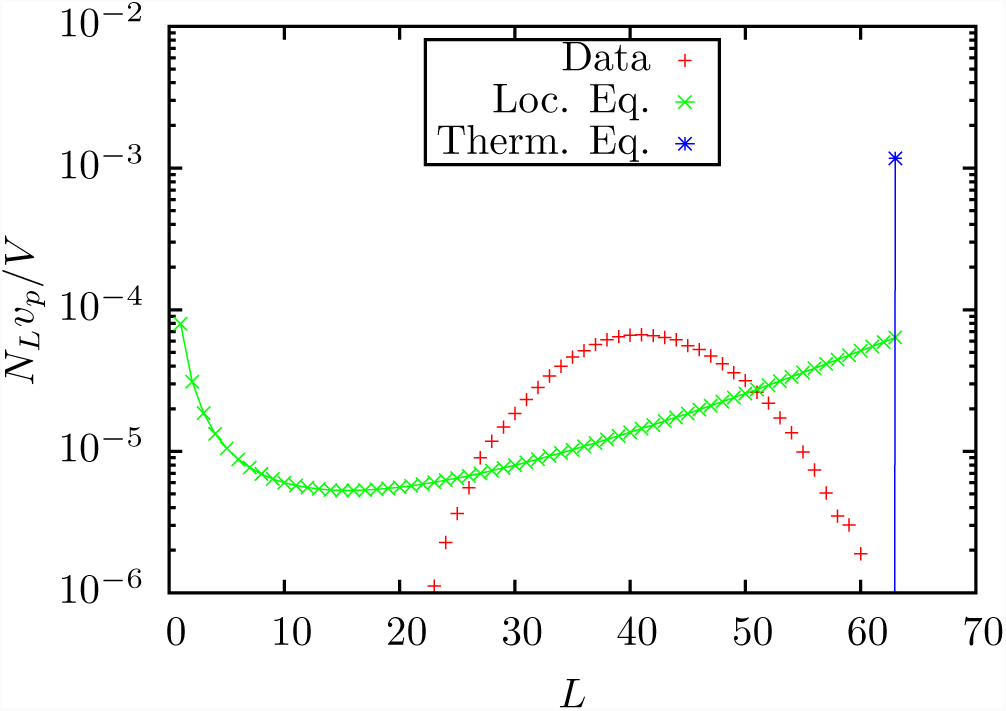
*N*_*L*_*v*_*p*_*/V* values for the copolymer system for which data are given in Figure 6-13c of [18] and corresponding equilibrium distributions. *R*_*L*_ = 0.14 and *R*_*T*_ = 0.72.

**FIG. 11:**
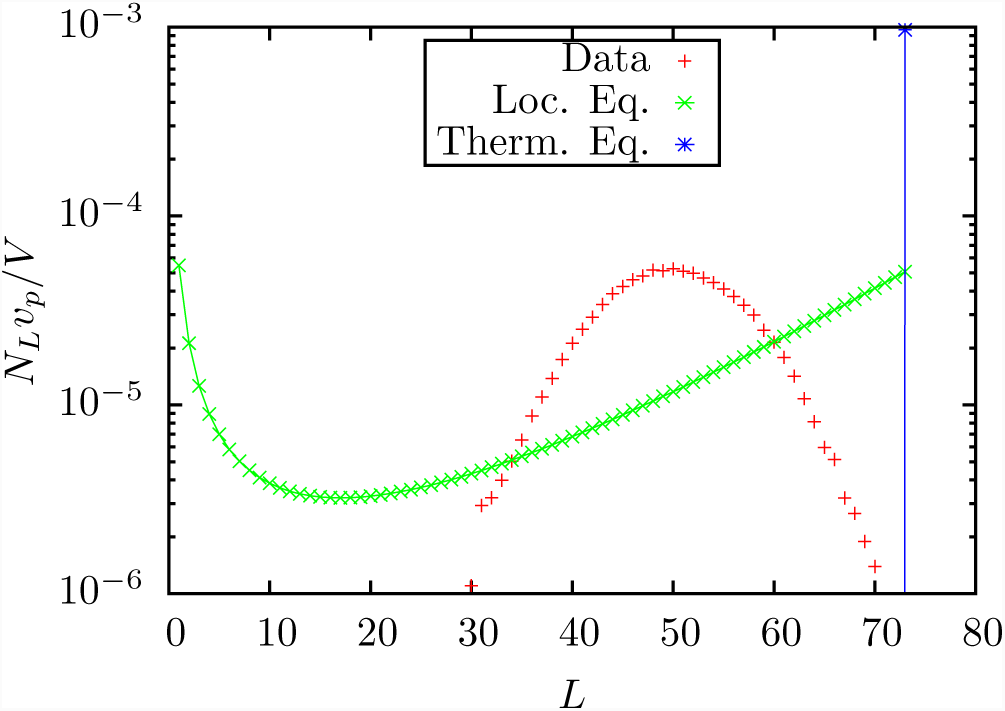
*N*_*L*_*v*_*p*_*/V* values for the copolymer system taken for which data are given in Figure 6-13d of [18] and corresponding equilibrium distributions. *R*_*L*_ = 0.14 and *R*_*T*_ = 0.72.

**FIG. 12:**
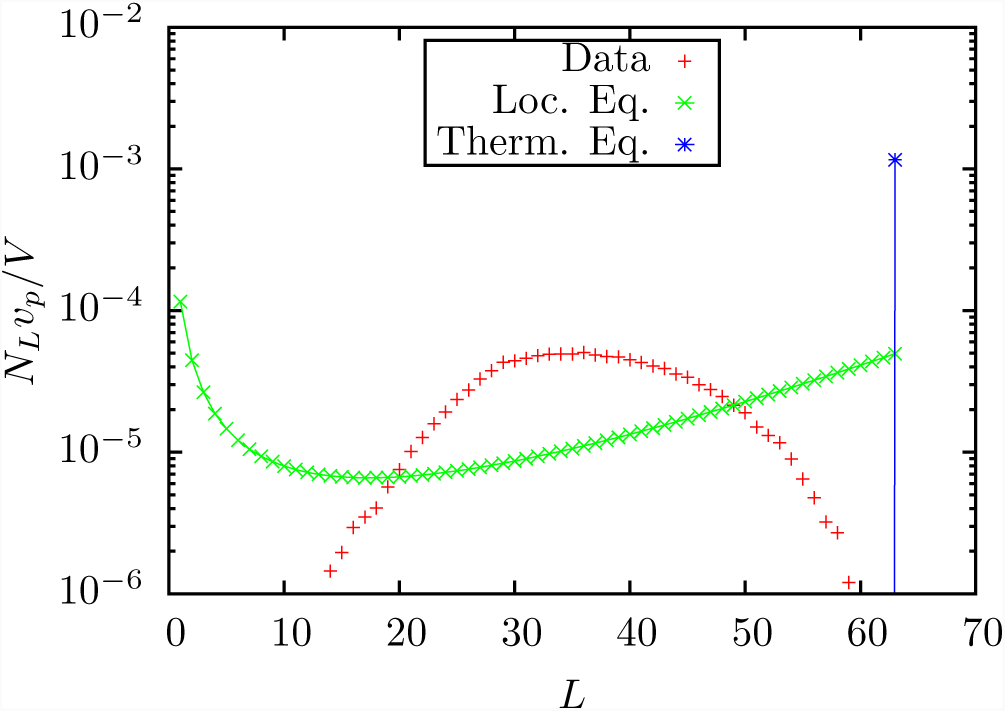
*N*_*L*_*v*_*p*_ */V* values for the copolymer system for which data are given in Figure 7-15 Left of [18]i and corresponding equilibrium distributions. *R*_*L*_ = 0.14 and *R*_*T*_ = 0.72.

**FIG. 13:**
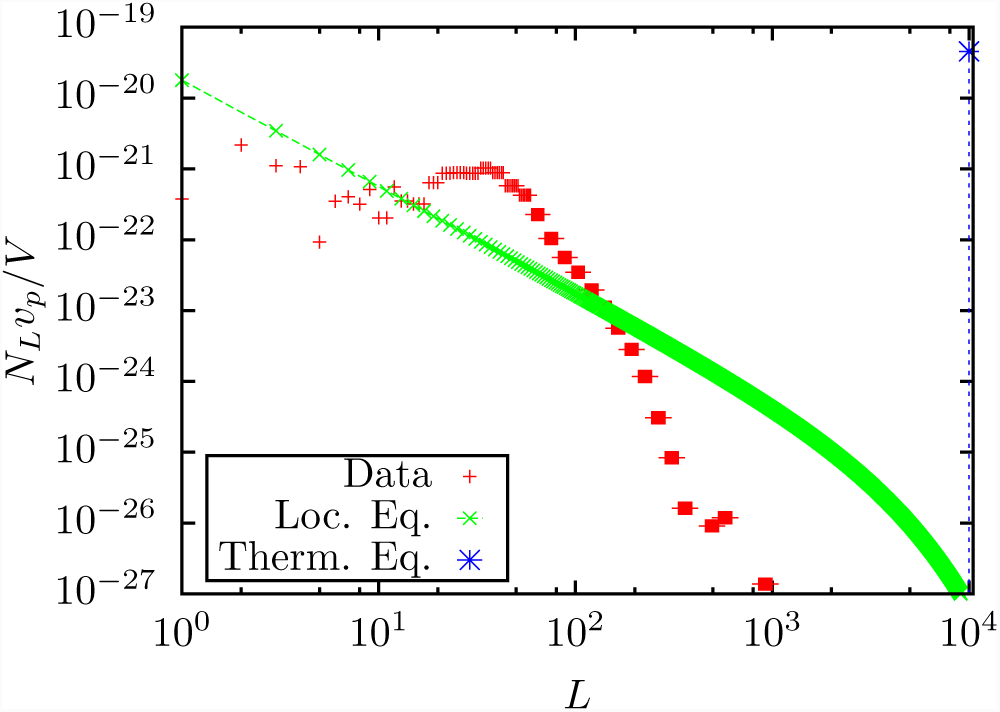
*N*_*L*_*v*_*p*_*/V* values for the Titan atmosphere at 1013km above the surface and corresponding equilibrium distributions.. *R*_*L*_ = 0.29 and *R*_*T*_ = 0.71.

**FIG. 14:**
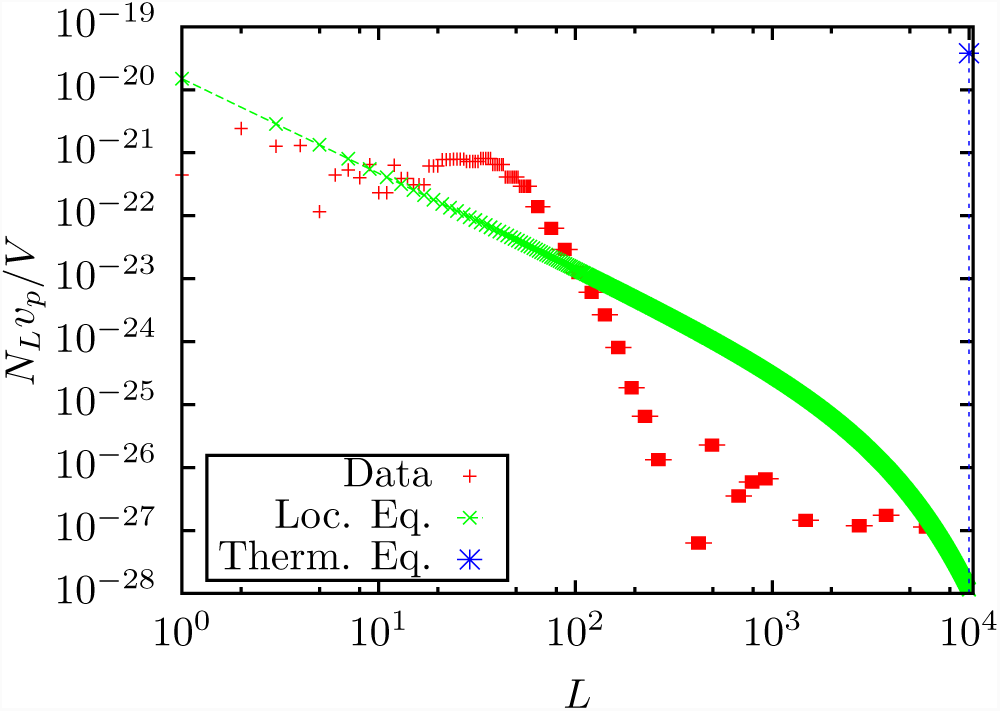
*N*_*L*_*v*_*p*_*/V* values for the Titan atmosphere at 1032km above the surface and corresponding equilibrium distributions. *R*_*L*_ = 0.29 and *R*_*T*_ = 0.71.

**FIG. 15:**
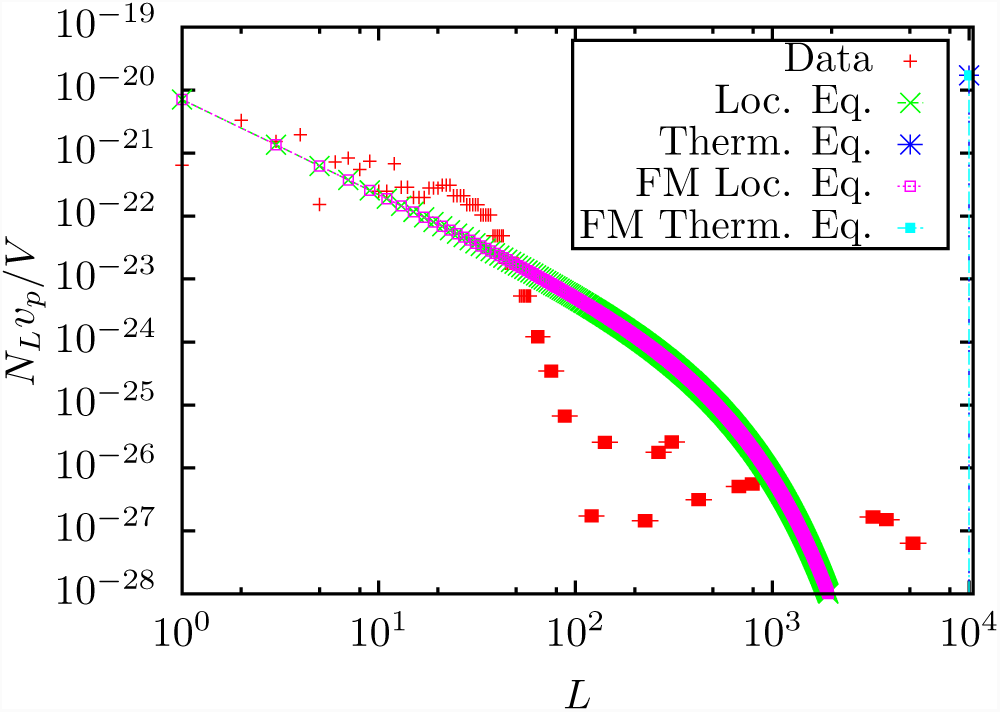
*N*_*L*_*v*_*p*_*/V* values for the Titan atmosphere at 1078km above the surface and corresponding equilibrium distributions. *R*_*L*_ = 0.28 and *R*_*T*_ = 0.73. The curves labeled ‘FM Loc. Eq.’ and ‘FM Therm. Eq.’ are the local and thermal equilibrium, respectively, calculated using the full model as described in [9].

**FIG. 16:**
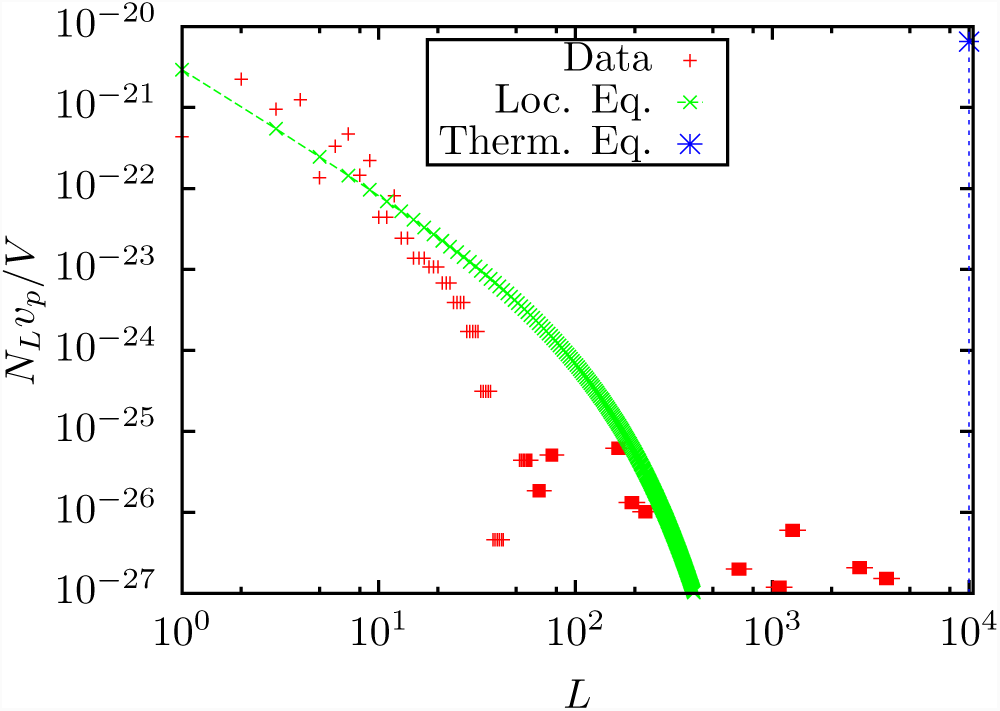
*N*_*L*_*v*_*P*_ */V* values for the Titan atmosphere at 1148km above the surface and corresponding equilibrium distributions. *R*_*L*_ = 0.32 and *R*_*T*_ = 0.77.

**FIG. 17:**
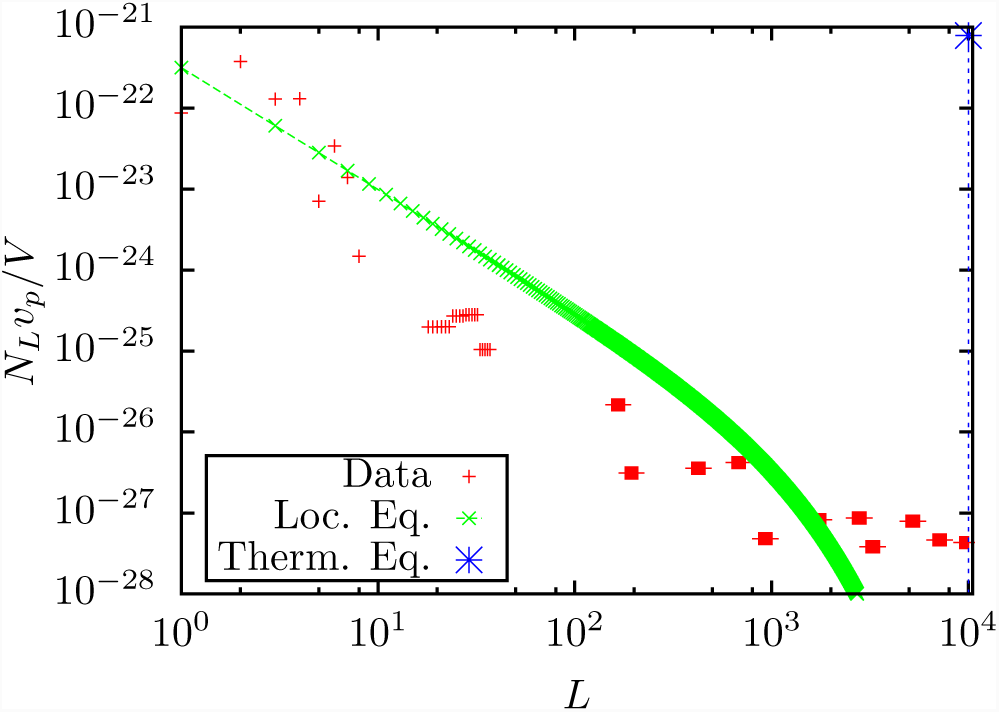
*N*_*L*_*v*_*p*_*/V* values for the Titan atmosphere at 1244km above the surface and corresponding equilibrium distributions. *R*_*L*_ = 0.33 and *R*_*T*_ = 0.81.

**TABLE IV:**
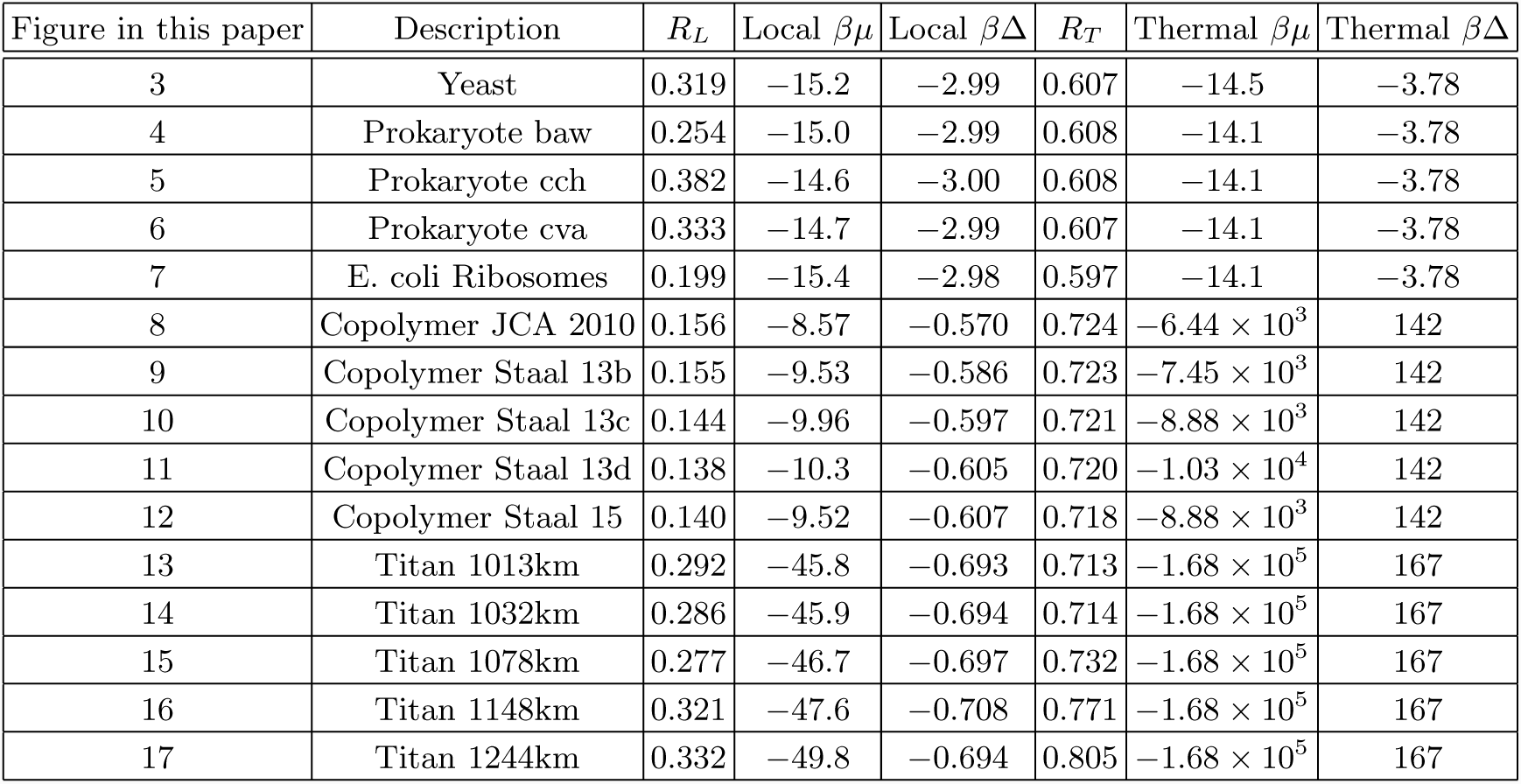
Values of parameters found here characterising the observed population distributions of the systems considered. (Equations 7 and 6 were solved with six figure precision but only three significant figures are reported here.)

In figure 18 we show a scatter plot of the average polymer length versus the standard deviation of the length distribution for the same set of systems. The next figure 19 shows the values of *β*Δ and *βμ* found for the isolated equilibrium state in each case.

**FIG. 18:**
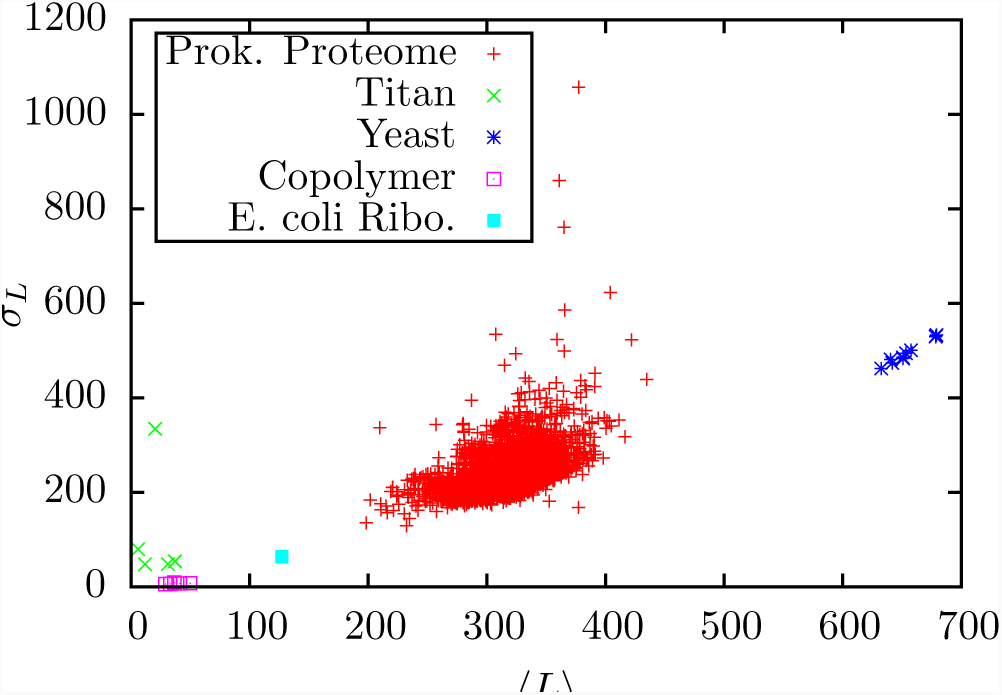
Average length versus standard deviation of the length (*σ_L_*).

**FIG. 19:**
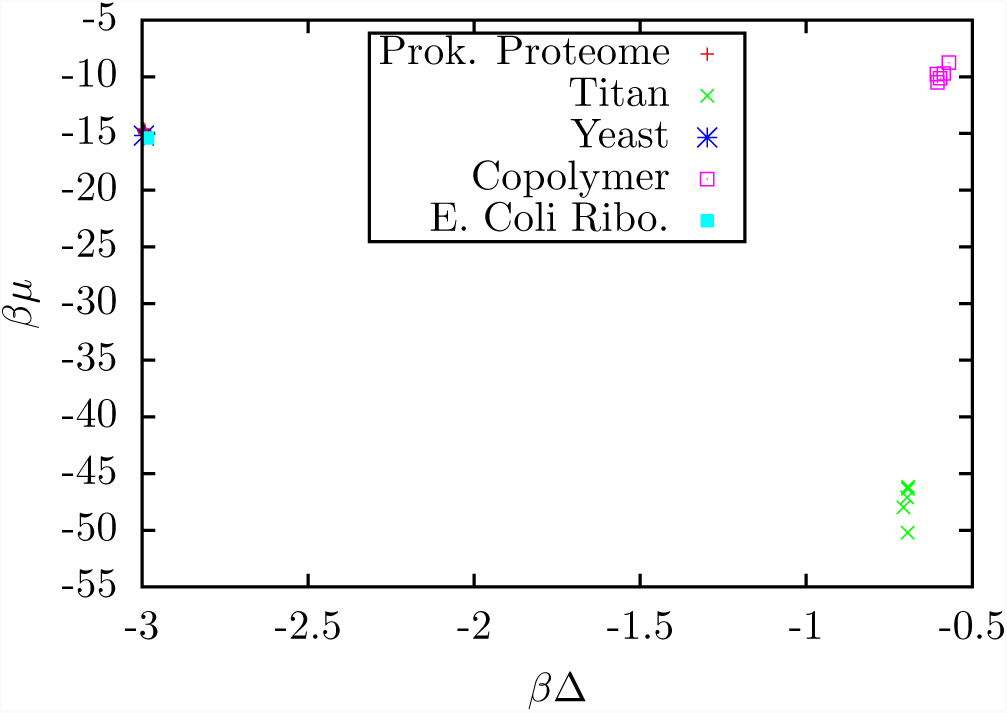
Scatter plot of *βμ* and *β*Δ for the various systems.

## V. DISCUSSION AND CONCLUSIONS

The results show quite definitely that the prokaryotic and yeast proteomes are far from chemical equilibrium with the ambient environment as measured by nearly identical values of about *R*_*T*_ ≈ 0.61 found for all of them. The reason for this is that the large number (b=20) of available monomers (amino acids) plays a significant role in determining the thermal equilibrium population distribution. (For all cases considered here, the terms −1 in the numerator and denominator of equation 2 can be ignored (the ‘Gibbs limit’).) The dominant factor determining the equilibrium distribution is *e*^(ln^ ^*b*+*β*Δ)*L*^. When Δ < 0 as it is for polypeptides the exponent can be negative. If the exponent is positive the thermal equilibrium is predominantly long polymers whereas if it is negative, the distribution is predominantly short polymers. This is a manifestation of the competition between entropy, which in the case of peptides drives the system toward long polymers, and energy, which drives it toward monomers. When *b* = 20 the external temperature, (or the bond energy) must be finely tuned to avoid one of these extremes and for most reasonable values of Δ and *β* the fine tuning does not occur and the thermal equilibrium state is described by one of the extremes. The *R*_*T*_ value is then the distance in the space of coarse grained populations between one of those extreme cases (all monomers or all polymers of maximum length) and the actual, nonequilibrium population distribution. Because the actual polymer distributions of the living systems are all similar and the thermal equilibrium distribution is at one of the extremes, the *R*_*T*_ values tend to be the same. However because of the sensitivity of the thermal equilibrium population to the value of *β*Δ one finds one or the other of the two extremes with values of Δ which are within the range of measured peptide bond energies. We illustrate this in Figure 20 where we show data for a prokaryote analysed using the average (−2.2 kcal/mol) of the values of the peptide bond energy reported in reference [8] and the results of the same analysis using the average plus one standard deviation and the average minus one deviation. For the least negative value of *β*Δ the thermal distribution jumps from all monomers to all polymers of maximum length.

**FIG. 20:**
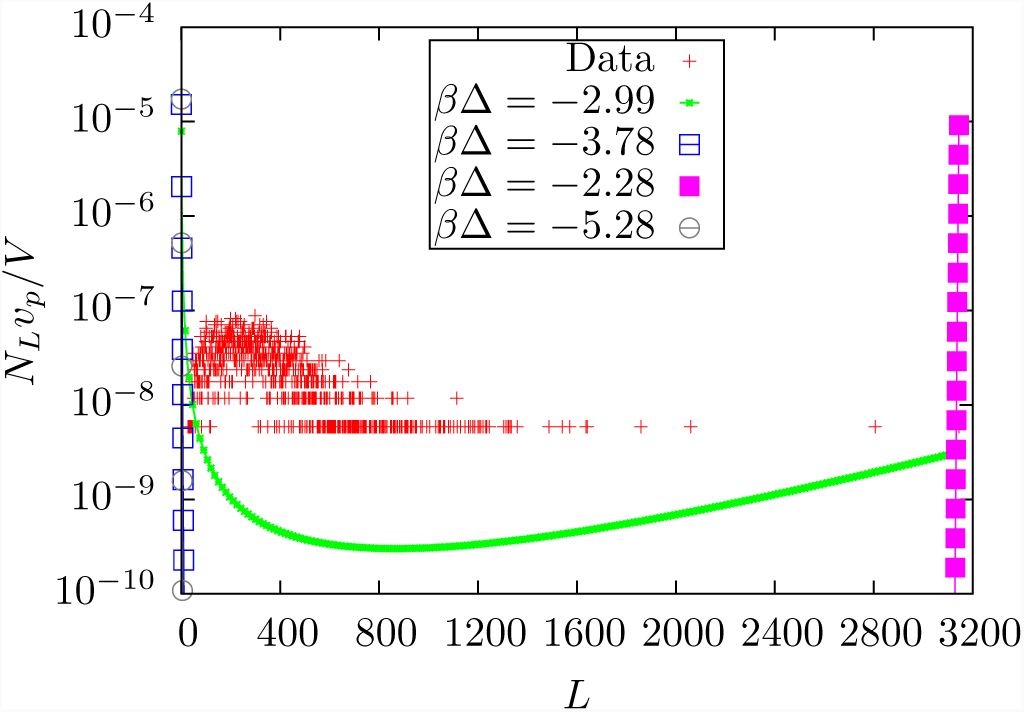
The data of figure 6 compared to equilibria at different values of *β*Δ. *β*Δ = −2.99 corresponds to the local equilibrated distribution, *β*Δ = −3.78 is the thermally equilibrated distribution using a bond energy of −2.2 kcal/mol and a temperature of 293K, *β*Δ = −2.28 is the distribution using a bond energy of (−2.2 + 0.875) kcal/mol and a temperature of 293K, and *β*Δ = −5.28 is the distribution using a energy of (−2.2 − 0.875) kcal/mol and a temperature of 293K. Here −2.2 kcal/mol and 0.875 kcal/mol are the average and standard deviation respectively of the protein bond energies found in [8].

We conclude that within our model, with ambient temperatures in the range of 100’s of degrees, a system of polypeptides with a large *b* (which is 20 for terrestrial biochemistry) will be very sensitive to the external temperature and will be driven toward long polymers at higher temperatures and monomers at lower ones with a sharp transition in between. The exact transition will be determined by the value of Δ (< 0). In the real systems, that value of the peptide bond energy varies from one amino acid pair to another and the corresponding sharp transition will be smeared by the distribution of bond energies. We illustrate these points in another way in Figure 21 where we show the calculated value of *R*_*T*_ for a series of *β*Δ values for one of the data sets. There is a sharp minimum in *R*_*T*_ when the thermal distribution is at the tipping point between all monomers and maximum length polymers. The minimum is close to but different from the local value of *β*Δ which was calculated from the protein energy and polymer number

**FIG. 21:**
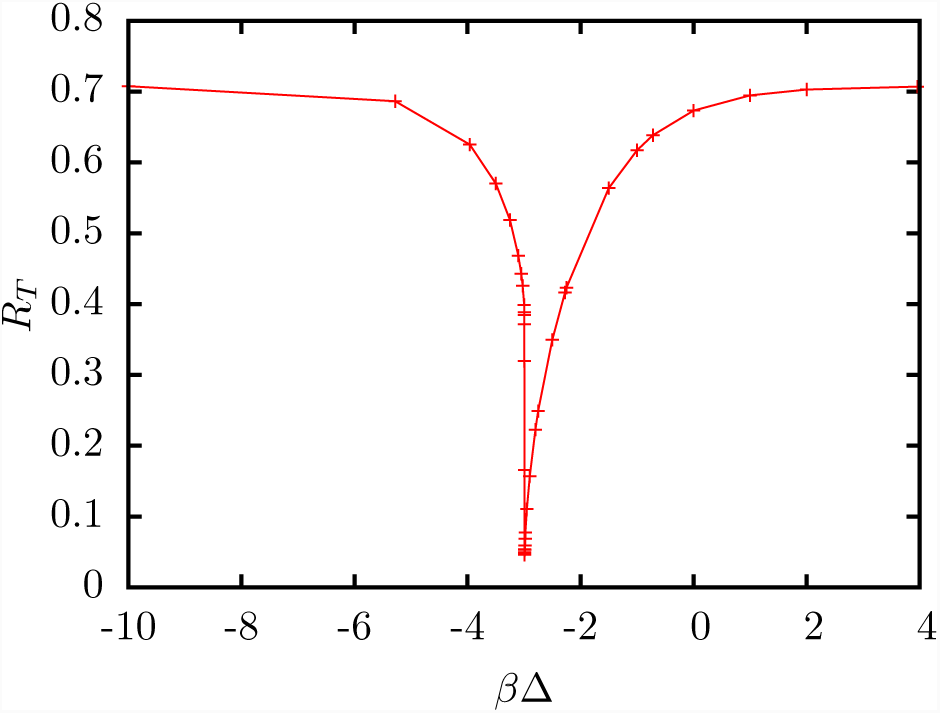
*R*_*T*_ as a function of *β*Δ for a fixed data set..

The implication seems to be that the observed protein distributions in the living systems are quite close to a local distribution which is quite unlikely since it requires a fine tuned value of *β*Δ to be produced in a thermal environment. One may construct the following plausibility argument to speculatively explain this: Generally, a large number of possible chemical configurations must be explored in order to find those special ones which result in lifelike properties such as autocatalysis. If a system is in a thermal environment which strongly favors all monomers or all long polymers, not many configurations will be explored. Hence the systems from which a lifelike system is most likely to emerge will be those which are thermally fine tuned to be in the intermediate region in which the system has a spread of polymer lengths and, we expect, large fluctuations in the polymer length distribution. Once autocatalysis begins, the system may become self stabilizing and its local distribution will differ from the one imposed by its thermal environment, as we observe for the prokaryotes, the yeast and the ribosome. Hence we suggest that the ‘local’ chemical equilibrium associated with the polymer density and bond energy density of these living systems, which is quite close to the actual population distributions (though the values of *R*_*L*_ which measure this disequilibrium vary between about 0.25 and about 0.38) might be a relic of the early thermal environment in which those living systems originally evolved. The needed value of temperature is easily estimated from the relation ln *b* = −*β*Δ from which *T* = |Δ| */*(*k*_*B*_ ln *b*). With the average bond energy we have used and *b* = 20 this is within a few degrees of the boiling point of water at 1 atmosphere.

These speculations suggest experiments with peptide systems in which the value of the effective *β*Δ is varied over a very fine scale to find the tipping point and discover if any interesting behavior emerges near it. For a more careful estimate of the needed temperature one could take the expected (or measured) temperature dependence of the bond energy into account and solve the resulting implicit equation for *T*.

The living systems have values of the parameters *R*_*T*_ and *R*_*L*_ which are robustly distinct from those for the copolymer (*R*_*T*_ ≈ 0.725, *R*_*L*_ ≈ 0.15) and the atmosphere of Titan 0.71 < *R*_*T*_ < 0.82, 0.28 < *R*_*L*_ < 0.33. In these two measures, Titan is mainly distinguished from the living systems by the values of *R*_*T*_. Its values of *R*_*L*_ fall within the same range as those of the living systems. Of course many other features of the upper atmosphere of Titan differ from those of the living systems. We regard it as a strength of the present method of analysis that these dimensionless measures of disequilibrium permit a dimensionless quantitative comparison of such dissimilar systems with regard to this feature.

Note that the copolymer systems are far from equilibrium by design: Most combinatorial combinations are excluded by the preparation procedure which assures that the molecules will all consist of a string of one type of monomer attached by one bond to a string of the other type of monomer whereas the corresponding approximate equilibria which we find take account of the possibility of mixing the two types of monomers in all possible ways along the chain.

All the molecules detected in the Titan atmosphere are almost certainly not linear as we have assumed in calculating the equilibrium distributions. Bonding rules assure that the energies will not be greatly affected by that error, but the degeneracies *G*_*L*_ which describe the numbers of molecules containing *L* monomers may be, particularly for small *L*. If as we anticipate, the error is larger for smaller *L* then this will shift the equilibrium distributions toward small *L* and it appears that this might increase the values of *R*_*L*_. The analysis could be repeated with more detailed models of the small molecule chemistry of the Titan atmosphere, but the idea here was to use only the empirical data available, and that data does not currently provide the information on the distribution of molecular types which is required.

There are some issues with the *R*_*T*_ values for the Titan atmosphere: We used a reported value [12] for the temperature of the Titan atmosphere in the analysis. But that is not really consistent because we are finding that the Titan atmosphere is not in equilibrium and therefore cannot itself be characterized by a temperature. Secondly the temperature used for the Titan atmosphere in the determination of *R*_*T*_ is much lower (*T* = 120*K*) than the characteristic terrestrial temperature (*T* = 293*K*) used in the corresponding analysis of the living systems. However we have recalculated *R*_*T*_ for all the Titan data assuming an ambient temperature of 293 K and find negligible changes.

We note that the molecular density of the organic molecules in the Titan upper atmosphere, both in absolute terms (particles per unit volume) and in dimensionless form as reported in figures 13-17, is much less than that of the living systems. One point which we are making in this work is that we can nevertheless compare its degree of equilibration with that for the living systems by a common measure. It has been a primary goal of our program to find such measures, which can be used to analyse and compare systems found elsewhere to determine whether they have lifelike properties without imposing excessive terrocentric bias. We do not have data on the atmosphere of Titan near its surface, where the pressure exceeds that of the earth’s surface atmosphere. However, as discussed briefly below, we do not anticipate the chemistry on the surface to be as far from equilibrium (and thus to have such large values of *R*_*T*_ and *R*_*L*_) as that in the upper atmosphere, unless of course there is actually a biosphere on or below the solid surface of Titan.

The selection of the Titan data as a test case for applying these measures is based in part on our earlier work on Kauffman models, which suggest that very dilute chemical systems might have a better chance than denser ones of not falling into chemical equilibrium during their temporal evolution and this appears to be a minimal requirement for the spontaneous development of lifelike molecular dynamics. It has been known for a long time [22] that chemistry in the dilute environment of space, both in the upper atmosphere of earth and other planets and in the interplanetary medium, does in fact avoid falling into chemical equilibrium while sustaining substantial chemical activity. That condition is much closer to that of the small *p* Kauffman models that we previously simulated and which led to nonequilibrium, possibly lifelike, dynamics, than to the environments of ponds and oceanic trenches which are often speculated to be possible sites of prebiotic evolution. Astrophysical energy sources for sustaining prebiotic chemistry in such environments have been extensively studied (eg)[23]) and many of the chemical constituents believed to be required are present (eg [4, 24]).

## Acknowledgments

This work was supported by the United States National Aeronautics and Space Administration (NASA) through grant NNX14AQ05G and used the used the computational resources of the Minnesota Supercomputing Institute, the Open Science Grid, the University of Minnesota School of Physics and Astronomy Condor cluster, and the NASA Advanced Supercomputing division Pleiades supercomputer. We thank Aaron Wynveen for helpful discussions and Ravindra Desai, Joao Paulo, Bastiaan Staal and Gabriel Vivo Truyols for answering questions about their work and supplying us with their data. The Titan data is available on NASA’s Planetary Database System as well, in summary form in [2]

## Appendix A: Conversion of Titan data

The reported data is of the form (log *L*_*i*_, *yL_i_*), where *yL_i_* is the number of counts in bin *L*_*i*_ centered at log *L*_*i*_ and *L* is the number of carbon or nitrogen atoms in a molecule. (The Cassini mass spectrometer did not have sufficient resolution to distinguish carbon from nitrogen masses.) To convert the reported distributions to a distribution as a function of *L* we suppose that the number 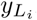 in the 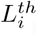 bin represents a uniform dis *I* tribution over the range (1*/*2)(log(*L*_*i* − 1_) + log(*L*_*i*_)) and (1*/*2)(log(*L*_*i*+1_) + log(*L*_*i*_)). On a linear scale the corresponding range for *L* is from 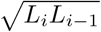 to 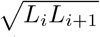. We round the lower limit to the nearest larger integer and the upper limit to the nearest lower integer and attribute a count value of 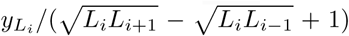 to each integer within the interval.

## Appendix B: Bond energies for Titan data

Because the N-N bond energy is much lower than the C-N and C-C bond energy as reviewed in Table II, the model for the energy expressed in the relation 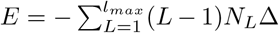; cannot be used and the calculation of the equilibrium distributions involves, in principle, a more complicated statistical mechanical calculation which is described elsewhere[9]. Here we describe a kind of mean field approximation in which we use an average bond strength 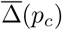 in the expression for the energy: 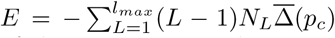where *p*_*c*_ is the fraction of the monomers in the system which are carbon. The average value which we use is

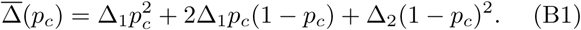

where Δ_1_ is the bond energy of the C-C and C-N bonds (assumed to be the same) and Δ_2_ is the bond energy of the N-N bond. The expression implicitly assumed that the probability of finding a carbon at a site is *p*_*c*_ independent of its environment and that is not expected to be a very good assumption. To test the effects of the error, we consider the maximum and minimum energies per bond which a polymer of long length *L* could have given *p*_*c*_ and show that the results would not be much affected by using these extremal values.

To find the average bond strength which gives the minimum energy consider a polymer of length *L* and *L* − 1 total bonds and that *L*_*c*_ = *p*_*c*_*L* of the monomers are carbon. The answer depends on whether *L*_*c*_ ≤ *L/*2 or not. In the former case, the total energy is minimized by starting with a nitrogen atom and then alternating between carbon and nitrogen until all of the carbon atoms are used giving a sequence *NCNCNCNCNCN N*. (The NCNCNCNCNCN sequence can be placed anywhere in the chain, giving a degeneracy, but we do not consider that here.) For each carbon atom there are two carbon-nitrogen bonds, the rest are nitrogen-nitrogen bonds. This gives a total energy of −2Δ_1_*L*_*c*_ − Δ_2_(*L* − 1 − 2*L*_*c*_). For the other case *L*_*c*_ *> L/*2 start with all C atoms replace less than half of them with N atoms. This can be done without introducing any N-N bonds and, since we assume that the C-N and C-C bond energies are equal we have a total minimum energy of −(*L* − 1)Δ_1_. Dividing these energies by *L* and taking the large *L* limit with *p*_*c*_ = *L*_*c*_*/L* fixed we obtain an average bond energy giving a minimum polymer energy of

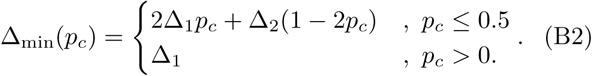

The calculation of the average bond energy which maximizes the total energy (Δ_max_) given *L* atoms is simpler: One must maximize the number of N-N bonds and that is done by connecting all the carbon atoms together, connecting all the nitrogen together, and then connecting the two with a single carbon-nitrogen bond (*C … CCNN … N*). The total energy is given by −Δ_1_*L*_*c*_ − Δ_2_(*L* − 1 − *L*_*c*_). Dividing this total energy by *L* and taking the limit *L* → ∞ yields the average bond energy which maximizes the total energy:

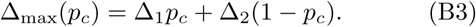

The three bond energies, 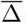, Δ_min_, and Δ_max_ are plotted in figure 22. Using each of these three bond energies for all the bonds we computed the values of *βμ,β*Δ for the Titan data for some test cases and found that the results were not sensitive to the value used. Using the three bond energies 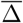, Δ_min_, and Δ_max_ for calculating the thermal equilibrium length distribution for the Titan 1078km data it was found that the value of *βμ* changed by only 1% and there was no discernible change in the value of *R*_*T*_. We also have preliminary, results plotted in 15, for the full model as described in [9] where one sees that the full model gives results quite close to those found by the approximate approach described above and used to obtain the results in the rest of the Titan data considered here.

**FIG. 22:**
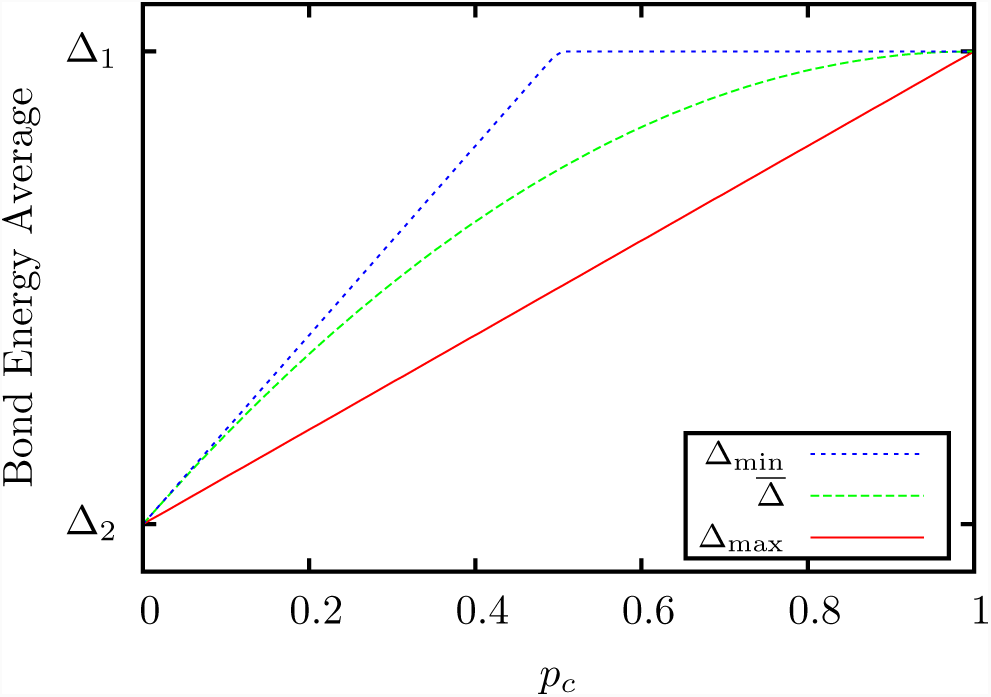
The three bond energies defined in Appendix B (equations (B1), (B2),(B3)) as a function of the carbon proportion *p_c_*.

